# Rationalizing Translation Elongation by Reinforcement Learning

**DOI:** 10.1101/463976

**Authors:** Hailin Hu, Xianggen Liu, An Xiao, Sen Song, Jianyang Zeng

## Abstract

Translation elongation plays a crucial role in multiple aspects of protein biogenesis. In this study, we develop a novel deep reinforcement learning based framework, named RiboRL, to model the distributions of ribosomes on transcripts. In particular, RiboRL employs a policy network (PolicyNet) to perform a context-dependent feature selection to facilitate the prediction of ribosome density. Extensive tests demonstrate that RiboRL can outperform other state-of-the-art methods in predicting ribosome densities. We also show that the reinforcement learning based strategy can generate more informative features for the prediction task when compared to other commonly used attribution methods in deep learning. Moreover, the in-depth analyses and a case study also indicate the potential applications of the RiboRL framework in generating meaningful biological insights regarding translation elongation dynamics. These results have established RiboRL as a useful computational tool to facilitate the studies of the underlying mechanisms of translational regulation.

## 1 Introduction

Translation elongation plays an essential role in protein biogenesis [1, 2] and has been increasingly recognized to associate with human diseases [3, 4]. However, despite the numerous research efforts in this field [5–14], the underlying mechanisms for the regulation of translation elongation still remain largely elusive.

In recent years, the accumulation of ribosome profiling data [15, 16] has provided an unprecedented opportunity for the applications of data-driven methods, especially machine learning approaches, in understanding the regulation of the translation elongation rate *in vivo*. In a typical ribosome profiling experiment, the mRNA fragments protected by ribosomes are captured and sequenced in a high-throughput manner, leading to a quantitative measurement of ribosome density at codon resolution. Based on these data, several computational models, e.g., RiboShape [17] and RUST [18], have been proposed to predict the distributions of ribosome footprints along the mRNA transcripts. More recently, Tunney et al. [19] have proposed a new approach called iχnos to predict ribosome density from an input sequence neighborhood through a feed forward neural network, which shows superior prediction performance over previous methods. However, few of these methods pay attention to the specific roles of different sequence features, e.g., codons, in a given mRNA context. Instead, they only provide a global measurement to evaluate the average effect of each feature on the local ribosome density.

In this study, we develop a deep reinforcement learning based framework, named RiboRL, for the accurate recovery of experimentally determined ribosome density and the rationalizing of translation elongation dynamics at codon level. Intuitively, we assume that the contribution of each sequence element, e.g., codon or nucleotide, to the local translation elongation rate is non-uniform and depends on a specific sequence context. Therefore, in contrast to the previous prediction methods which feed the whole sequence context into a universal regressor, RiboRL first performs a context-dependent feature selection procedure for each sequence input, aiming to select the most relevant sequence features, also called rationales, for this specific learning task. To mimic the biological translation process, we model feature selection as a sequential decision process, following the reading order of a ribosome. Based on the generated rationales, a predictor is subsequently trained to predict the ribosome density. As the data label does not provide direct supervision information on the importance of individual codon positions, the training process of RiboRL is conducted by reinforcement learning in a trial and error manner through the coordination of a designed reward system which explicitly optimizes both quality and sparsity of the selected rationales.

The advantage of introducing the context-dependent feature selection scheme into our deep learning framework is two-fold. On one hand, the sparsity constraint forces the model to select the most relevant sequence features for prediction and therefore boosts the model performance, which in fact applies the same philosophy as in current sparsity regularization based statistical learning approaches [20]. On the other hand, it also greatly enhances the explainability of our deep learning model, enabling the application of RiboRL for mining potentially intriguing biological insights into translation elongation dynamics.

In this work, extensive tests using currently available high-quality yeast ribosome profiling data have demonstrated that RiboRL can significantly outperform the current state-of-the-art method as well as other commonly used deep learning architectures in ribosome density prediction. Moreover, by comparing the quality of the features selected by different attribution methods, we have also shown that RiboRL is able to generate the most informative sequence features to facilitate the precise prediction of ribosome density. In addition, genome-wide analyses and a case study using RiboRL have also revealed several interesting biological findings, further establishing the biological relevance of our model. These results have demonstrated that RiboRL can serve as a powerful tool to explore the sequence features of translation elongation dynamics and analyze various translation elongation-related phenomena, which will provide useful insights into understanding the underlying mechanisms of protein biogenesis. RiboRL is available as an open source software at https://github.com/Liuxg16/RiboRL.

## 2 Methods

### 2.1 Ribosome profiling datasets

The ribosome profiling data of *S. cerevisiae* collected by Weinberg et al. [12] were retrieved under the accession number GSM1289257. Reads were first trimmed to remove 8-base randomized barcode sequences from the 5’ end and the linker sequence (TCGTATGCCGTCTTCTGCTTG) from the 3’ end. To remove reads originating from ribosomal RNAs (rRNAs) and noncoding RNAs (ncRNAs), trimmed reads longer than 10 bases were further mapped to yeast or human rRNA and ncRNA sequences with bowtie2 version 2.1.0 [21], respectively. Filtered reads were then mapped to the customized transcriptome provided by [19] using hisat2 version 2.0.4 [22, 23]. In particular, each annotated coding DNA sequence (CDS) was extended by 13 nt on the 5’ end and 10 nt on the 3’ end to facilitate the mapping of the ribosome footprints spanning across the CDS boundaries.

For the calculation of codon occupancies and the assignment of A sites, we only kept those uniquely mapped reads with a length of 28, 29 or 30. For reads of length 28 or 29, A sites were assigned to the codon at position +14, 15 or 16 from the reads start, corresponding to frames - 2, 0, or −1, respectively. Similarly, for reads of length 30, A sites were assigned to the codon at position +15, 16 or 17, corresponding to frames −2, 0, or −1, respectively. Following the previous protocol [19], the raw read count for each codon was normalized by the average ribosome density of the corresponding transcript. The normalized ribosome density serves as the data label for our regression task.

### 2.2 The RiboRL Framework

In this study, inspired by [24], we propose a deep reinforcement learning based framework, named RiboRL, to predict ribosome density by effiectively incorporating context information that shapes the translation elongation dynamics (Figure 1). To boost the performance of ribosome density prediction as well as increase the explainability of our deep learning model, we employ a context dependent feature selection procedure to obtain the most relevant sequence features, which are also called rationales, for the prediction task. In contrast to the widely used attention mechanism [25], the RiboRL framework performs a binary feature selection operation which is trained using reinforcement learning in a trial-and-error fashion. Generally speaking, the main workflow for the RiboRL framework can be divided into two deep neural networks, namely the PolicyNet and the Predictor.

- The PolicyNet is a policy network which defines the probability of selecting a codon as a rationale based on the current input. In particular, it consists of two recurrent neural network (RNN) layers and is built upon the one-hot encoding of the input codon sequence. At each input codon position, the PolicyNet takes action to decide whether to select the codon information there as a feature to predict ribosome density. At this end, all the selected codons constitute the rationale set and are fed to the Predictor for the ribosome density prediction, while the unselected codons are masked as zeros to remove the codon information at those positions.
- The Predictor is composed of a bidirectional recurrent neural network (BiRNN) [26] and a fully connected layer. It predicts the ribosome density based on the selected context-dependent features, i.e., the rationales previously generated by the PolicyNet.

Our framework follows the basic architecture of an actor-critic algorithm [27] in reinforcement learning, in which one component (i.e., the PolicyNet) is designed to generate action sequences and the other (i.e., the Predictor) produces rewards according to the previously generated action sequences to prompt superior actions in the next trails. During the training process, the Predictor can be optimized through the standard back-propagation strategy, while the parameters in the PolicyNet are learned by the reinforcement learning algorithm to maximize the expected reward. Here, those two components are jointly trained to minimize the mean square error (MSE) between the prediction and the experimentally measured ribosome density.

**Figure 1:**
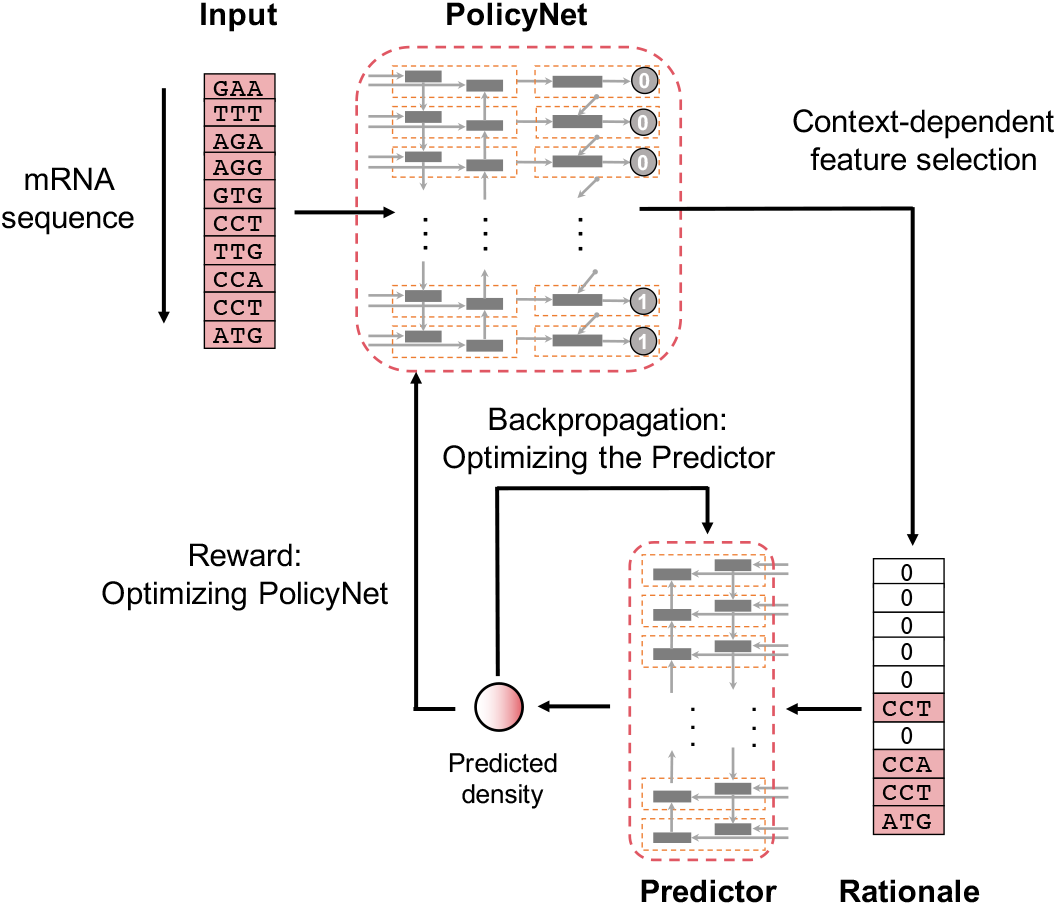
Schematic overview of the RiboRL pipeline, which mainly consists of two deep neural networks, i.e., the PolicyNet and the Predictor. See the main text for more details.

#### 2.2.1 Context dependent rationale generation by the PolicyNet

To provide an appropriate representation of the sequence context of translation dynamics, for each given codon at an A site (denoted as 0 position), we extract the codon sequence spanning from −5 to +4 positions as its input to our framework (Supplementary Figure S1). Then we denote the one-hot representation of the input codon sequence by ***X*** = (***x***_1_, ***x***_2_, …, ***x***_*T*_), where *T* stands for the number of input codons (which is set to 10 in this study). Let *D* denotes the total number of codon types, here we have each ***x***_*t*_ ∈ ℝ^*D*^ to encode the codon type information for a given sample. In particular, after indexing all the codon types, the *m*-th codon type is encoded as a binary vector of length of *D*, in which the *m*-th element is set to one while the others are set to zeros.

To capture the context-dependent information among the input codon sequence, we first use a bidirectional recurrent neural network (BiRNN) to extract features from the raw one-hot encoding representation of the input sequence. Formally, the BiRNN takes a sequence of codon features (***x***_1_, ***x***_2_, …, ***x***_*T*_) recurrently and then concatenates the two hidden states 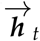 and 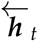. In other words, the hidden state ***h***̂_*t*_ corresponding to the time step *t* is defined as

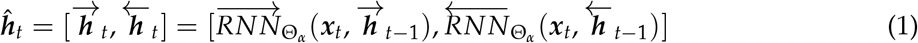

where 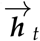 and 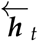 stand for the hidden states of the recurrent neural networks 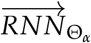 and 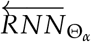 for opposite directions at the time step *t*, respectively, and Θ_*α*_ denotes the set of parameters in the neural networks. Here, we use the gated recurrent unit (GRU) [28] as our recurrent unit in BiRNN. Thus, the outputs of the GRU ***h***_*t*_ for each direction are computed by

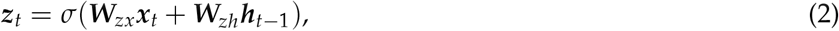

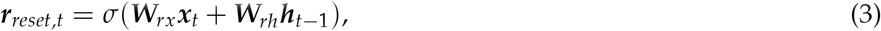

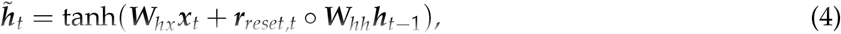

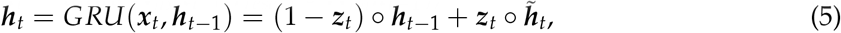

where ○ denotes element-wise product, and *σ* denotes the sigmoid function, ***W*** ’s stand for the weights, ***z***_*t*_ and ***r***_*reset*,*t*_ stand for update and reset gates, respectively, and the bias terms are omitted in the above equations for clarity.

Based on the feature representation generated by BiRNN, we employ another variant of RNN, (i.e., RNN with feedback) to generate actions, indicating whether to select a codon as a rationale. Rationale selection for a given sequence ***X*** can be equivalently defined as a series of binary indicator variables (*s*_1_, *s*_2_, …, *s_T_*), where each *s_t_* ∈ {0, 1} indicates whether codon *x_t_* is selected or not. We assume the rationale generation process, which can also be called action sequence in reinforcement learning, possesses the Markov property [29], i.e., the determination of the next action only depends on the current input and the complete previous history. Formally, the probability *P*(***s|X***) for generating context-dependent rationales ***s*** is given by

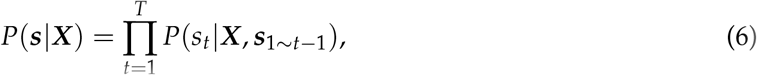

where ***s***_1*~t−*1_ means the variables from *s*_1_ to *s_t−_*_1_.

Specifically, to model the above conditional dependency relationship, the RNN with feedback considers the current input ***h***̂_*t*_, the previous state ***υ***_*t−*1_ and the previous actions ***s***_1*~t−*1_ together, which maintains dependency over the entire history and makes a decision at the current step. In particular, to capture the information from previous actions, we define the summation function *f* (·) = ∑ *s_i_* to represent the overall state of the previous historical actions. Note that the result of this summation is one-hot encoded as a ten-dimension vector and concatenated with the codon feature ***h***̂_*t*_ as the input to the RNN with feedback. Formally, the decision *s_t_*, after processing the *t*-th codon, is given by the policy distribution obtained through the following operations

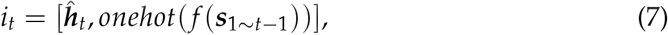

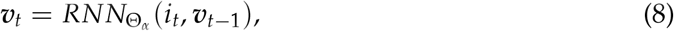

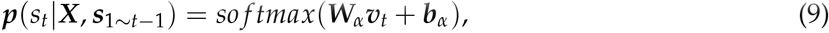

where ***W_α_*** and ***b_α_*** stand for the weights and the bias term, respectively. Note that as in widely used deep reinforcement learning frameworks [29, 30], during training we sample an action from its predicted distribution using *ε*-greedy methods [29] (Supplementary Notes), while for testing, we choose the action for the current step according to the maximum a posteriori probability, i.e., *s_t_* = *argmax* ***p***(*s_t_ |****X***, ***s***_1*~t−*1_). In this way, through combining the actions determined by the PolicyNet and the input codon sequence, we can obtain the rationale matrix ***R***, that is,

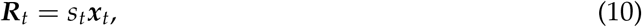

where ***R*** represents the processed one-hot representation of the raw input, in which the selected codons corresponding to the rationales maintain the original input information and those non-rationale codons are masked as zero and excluded for the downstream prediction.

#### 2.2.2 Predicting ribosome density with rationales

The Predictor models the distribution of ribosomes along the transcripts as a regression function of the rationales generated by the PolicyNet. The Predictor possesses two submodules. First, a BiRNN is employed to learn meaningful interdependencies and patterns from the input rationale matrix ***R***. Here, inspired by [31], we perform a max-pooling operation to select the maximum value for each dimension of hidden states over all the time steps to perform dimension reduction. Then, on the top of the max-pooling layer, a fully connected layer is employed to predict the ribosome density based on the extracted hidden features ***u***. Therefore, the detailed computations of the predicted ribosome density, denoted by *Predictor*_Θ*β*_ (***R***) can be expressed as

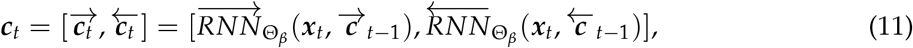

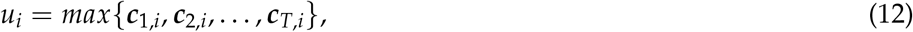

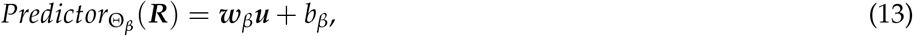

where ***c***_*t*_ stands for the vector of hidden states corresponding to codon *t* output by the BiRNN (i.e., 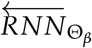 and 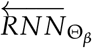), ***w***_*β*_ and *b_β_* stand for the weights and the bias term for the fully connected layer, respectively, and Θ_*β*_denotes the set of parameters in the Predictor.

#### 2.2.3 Learning

As described in the above subsection, the role of the Predictor is to predict the distribution of ribosomes as precise as possible. Therefore, it can be directly optimized by minimizing the expected mean square error over the distribution of the selected rationales, i.e.,

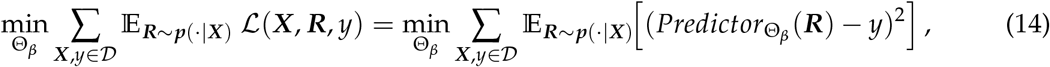

where *y* represents the true ribosome density, 𝒟 stands for the collection of training instances, ℒ stands for the loss function and Θ_*β*_ stands for the set of parameters in the Predictor.

On the other hand, the training of the PolicyNet is challenged by the lack of direct supervision information on the importance of individual codon positions from the training samples, and unfortunately the binary selection of features truncates the back-propagation of gradients [32]. Therefore, we apply a reinforcement learning strategy to train the PolicyNet, where the performance of rationale selection can be tuned by the well designed reward. Intuitively, we seek for a reward to encourage the PolicyNet to generate sparse and high-quality rationales, so that the Predictor can precisely and robustly output the true ribosome density based on the given rationales which are expected to represent the most relevant biological determinants of translation elongation dynamics. In particular, when we define the reward for the actions generated by the PolicyNet, in addition to considering the mean square error, we also use the *L*1 norm to constrain the vector ***s*** for the sake of sparsity. More specifically, we define the final reward, i.e., the reward given to the model after an entire action sequence as

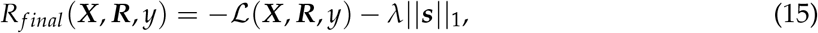

where *||* · *||*_1_ stands for the *L*1 norm, and *λ* stands for the penalty factor of sparsity. The training objective is to maximize the expected reward over sampled actions^1^, which can be expressed as

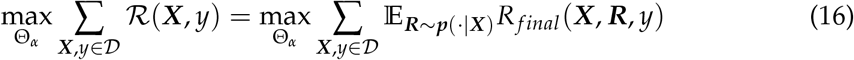

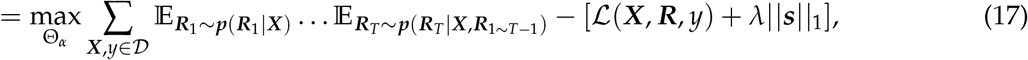

where *R*(***X***, *y*) represents the expected reward over all sampled actions and *R_final_* (***X***, ***R***, *y*) stands for the final reward given an action sequence ***s***.

During the training process, the optimization of the PolicyNet and the Predictor are mutually beneficial, i.e., the rationales generated from a better PolicyNet can promote the learning process of the Predictor, while a better Predictor will lead to give more accurate reward for the PolicyNet. In addition, their objectives are consistent in minimizing the mean square error. Therefore, we can jointly train these two modules at the same time, with a joint objective defined by

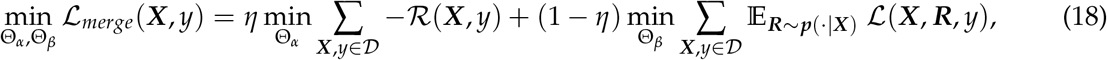

where *η* coordinates the learning speeds of the two modules. Here, we use the REINFORCE algorithm [33] to train the PolicyNet and the gradients with respect to the corresponding parameters are detailed in Supplementary Notes due to page limitation. To reduce the variance training results of REINFORCE, we subtract the reward by a baseline term (i.e., the average over *M* = 10 samples) and truncate the negative rewards as in [34, 35].

## 3 Results

### 3.1 RiboRL accurately predicts ribosome density

To evaluate the performance of RiboRL in predicting ribosome density, we first trained and tested our model using a high-quality yeast ribosome profiling dataset [12]. As in [19], to avoid the introduction of the potential noise caused by low sequencing coverage, the training of RiboRL was performed on a set of genes with high ribosome density. In particular, we selected a gene set with top 500 ribosome footprint density values and then split it into a training set with 334 genes (about 2/3) and a test set with 166 genes (about 1/3, denoted by the high-density test set), respectively.

We first compared our method with a recently published state-of-the-art method, namely iXnos [19], which adopted a feed-forward neural network to predict ribosome density and was shown to outperform the previous methods such as RUST [18] and Riboshape [17] (Table 1). Expectedly, with the advanced feature selection and modeling schemes, RiboRL achieved significant improvement in the regression performance of ribosome density, with an increase in Pearson correlation coefficient of 3.5% on this high-density test set. Note that this result was achieved using codon information as input alone. Similar to iXnos, we also noticed an increased prediction performance when adding nucleotide-level information as input (Supplementary Notes and Supplementary Table S1). For the sake of the explainability of our framework, in the subsequent analyses, we still used the RiboRL model with codon information as input alone.

**Table 1:** Comparison of prediction performance of RiboRL with that of different baseline methods in terms of Pearson correlation coefficient. The highest values are shown in bold. This comparison was conducted using codon sequence as input information.

| Model | Pearson’s $r$ (high-density test set) | Pearson’s $r$ (full gene test set) |
| --- | --- | --- |
| ixnos | 0.495 | 0.385 |
| CNN | 0.526 | 0.386 |
| BiGRU-Attention | 0.516 | 0.387 |
| BiGRU-Max-pooling | 0.527 | 0.370 |
| RiboRL | **0.530** | **0.397** |

In addition, we also showed that the RiboRL framework can outperform several other deep learning based approaches (Supplementary Notes), including the convolutional neural network [36], as well as the bidirectional gated recurrent neural network (BiGRU) with either max pooling (denoted by BiGRU-Max-pooling) or attention mechanism (denoted by BiGRU-Attention) (Table 1). The best hyperparameters of all methods were obtained through a grid search for best hyperparameters (Supplementary Table S2). Note that BiGRU-Max-pooling shares the same predictor with RiboRL, (which actually additionally leverages the PolicyNet for feature selection), while BiGRU-Attention depends on the soft attention mechanism to weigh and use the contribution of each input codon to obtain the final prediction. Notably, although all of them showed an increase in prediction performance, possibly by the introduction of a more sophisticated model architecture, RiboRL still outperformed all these methods, demonstrating the advantage of using the PolicyNet in our prediction framework.

As the aforementioned experiments were conducted mainly on transcripts with high ribosome densities, to ensure that the prediction performance can also be generalized to the majority of genes *in vivo*, we further tested our model on all the 5,114 genes that were detected in the ribosome profiling experiments [12] (denoted by the full gene test set) (Table 1). Consistent with previous reports, all the methods showed a decrease in prediction performance, mainly due to the decreased data quality caused by lower transcript abundance [19]. Interestingly, we found that on this dataset, RiboRL yielded significantly higher regression performance than other deep learning based methods, indicating the better generalizability of our model, especially when compared to BiGRU-Max-pooling. Overall, RiboRL showed a superior performance over all the other tested models, which greatly supported the contribution of reinforcement learning in effectively selecting the relevant features in predicting ribosome density.

### 3.2 RiboRL outperforms other methods in feature attribution

As the introduction of reinforcement learning-based feature selection significantly improved the prediction performance of ribosome density, we next tried to examine the quality of selected features, i.e., rationales, in our framework. We also compared our feature selection strategy with two families of commonly used attribution methods. The first family is the back-propagation-based methods, represented by integrated gradients [37] and DeepLift [38], which do not require the modification of the original network and only rely on the back-propagation procedure to pass the gradients or activation signals in a readily trained neural network. In comparison, the attention mechanism [25, 39–41] is implemented by adding an attention network (i.e., a neural network that determines the importance of each input position based on its associated features), which is trained simultaneously with the whole neural network to combine the feature representation of each position in the input codon sequence for the final prediction. The details of all the methods in this experiment can be found in Supplementary Notes and Supplementary Table 3.

Intuitively, an attribution method that is recognized to outperform another is expected to generate features that can yield better predictive power in the corresponding learning task. Therefore, in our experiment, we first selected the features generated by different attribution methods based on their importance scores assigned to individual input positions. The unselected positions were set as unknown, which was equivalent to removing the codon information in those positions. Subsequently, the features selected by different attribution methods were used to train, respectively, a bidirectional gated recurrent neural network as described in the Method Section to evaluate the prediction power of each set of the selected features. Similar results were also obtained using a simple multiple layer perceptron (MLP) regressor in this experiment (Supplementary Figure S2).

Expectedly, all the three attribution methods showed a great improvement over random selection, indicating their acquisition of important positions for predicting ribosome density. In particular, we found that for different feature lengths tested, the features selected by RiboRL provided the best prediction performance measured in terms of both mean square error and Pearson’s correlation coefficient (Figure 2). These results clearly demonstrated the success of our reinforcement learning-based strategy in selecting the most relevant features about ribosome density.

**Figure 2:**
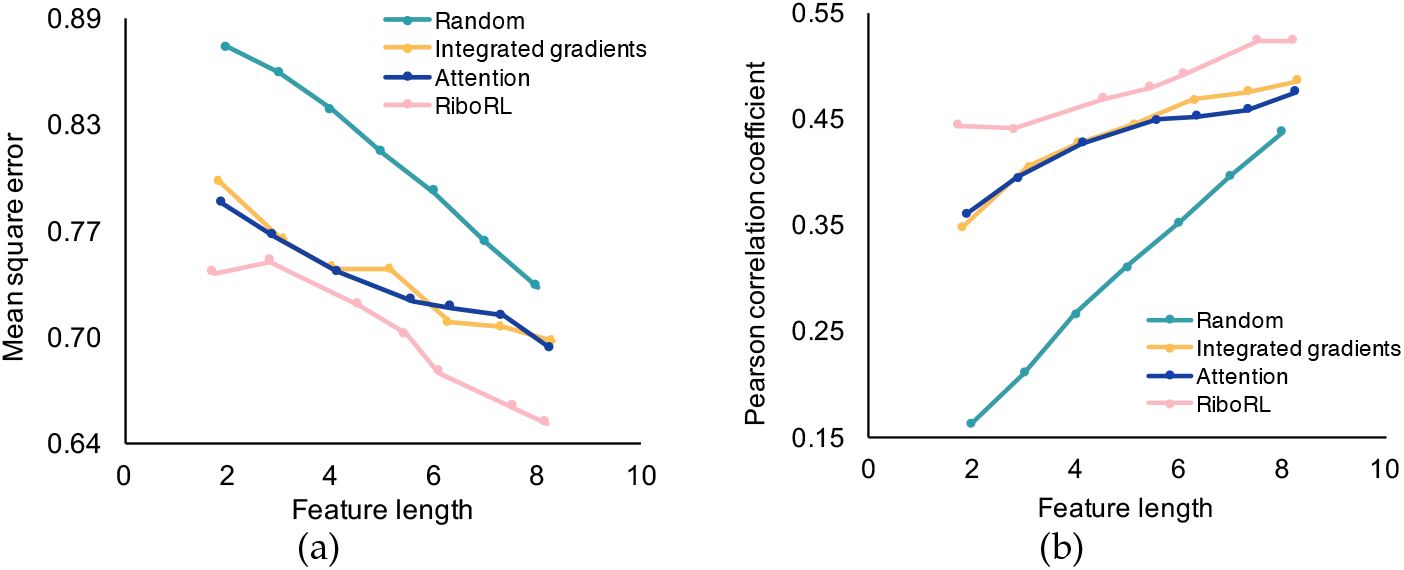
Comparison of different feature attribution methods in predicting ribosome density. Each codon in the input sequence context is scored by different methods to indicate the importance of that position. By adjusting the threshold of feature selection or the sparsity factor for RiboRL, features can be selected with different lengths and used to train a BiGRU regressor to evaluate their predictive power in terms of mean square error (a) and Pearson correlation coefficient (b).

### 3.3 RiboRL indicates the codon-level rules for translation elongation

Next, we attempted to analyze how RiboRL learns the biological rules that govern the regulation of translation elongation dynamics. Due to the sparsity constraint introduced in the feature selection stage (See Section 2.2.3), our final RiboRL model selected on average 75.2% of the input codons as rationales for ribosome density regression. Interestingly, we found that the fractions of being selected as rationales are associated with codon identity and display distinct distributions among the 61 amino-acid coding codons (Figure 3(a)). In particular, codons encoding charged amino acids and amino acids with special conformations, i.e., proline and glycine, are more frequently selected as rationales, which was consistent with previous reports [9, 12, 43]. In contrast, codons for hydrophobic amino acids are less involved in the final prediction, demonstrating the effects of amino acid species on rationale selection. We also noticed that the extreme rare codon encoding arginine (CGG) was 100% selected as rationale by the PolicyNet, indicating a particular role of this codon in translation elongation dynamics.

**Figure 3:**
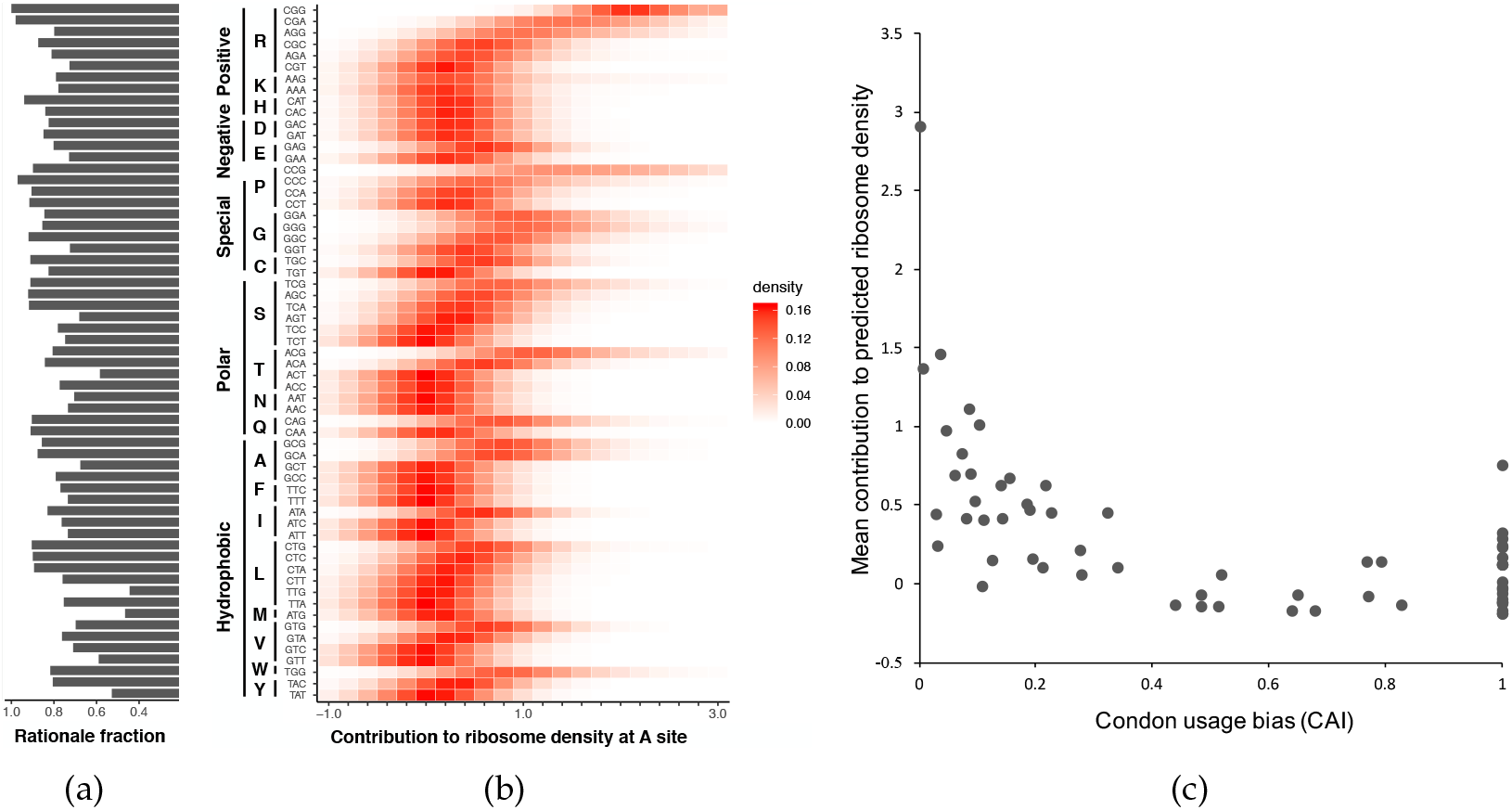
The effect of each codon species on translation elongation dynamics. The analysis was performed on the high-density test set. (a) The fraction of being selected as rationales by RiboRL for each codon species. (b) The contribution of each codon species to the predicted ribosome density at A site (smoothened by kernel density estimation [42]). The corresponding amino acids and their biochemical categories are also provided. (c) The mean contributions of individual codon species to the predicted ribosome density are associated with codon rarity. The codon usage is measured by the codon adaptation index (CAI).

To better characterize the contribution of each codon species to predicted ribosome density, we further performed a quantitative analysis in which we masked one codon in input information for the Predictor and calculated the difference in the predicted ribosome density with and without this specific codon information (Figure 3(b)). Here, the predicted ribosome density with masked input served as a reference score and the increase beyond this score thus represented the contribution of this masked codon. We mainly focused this analysis on the role of each codon at A site, as this site had been shown to act as the most important site determining the translation speed [12]. Note that as the non-rationale codons are already masked as zero in the input to the Predictor, our prediction was performed with only the codons corresponding to the selected rationales, which are expected to provide more biologically relevant information. Expectedly, we noticed a dramatic contribution of CGG (arginine), CGA (arginine) and CCG (proline) to the accumulation of ribosome density, presumably due to their amino acid properties and rarity among the synonymous codons [19]. Inspired by this observation, we further explored the correlation between codon rarity and the mean contribution to ribosome density at A site (Figure 3(c)), which would possibly explain the discrepancy in the effect of different synonymous codons of the same amino acid. Intriguingly, we found that the mean contribution of each codon species was negatively correlated with its codon usage (Pearson correlation coefficient −0.58), measured by the codon adaptation index (CAI) [44]. These results demonstrated that RiboRL is able to capture the biological determinants of translation elongation dynamics at codon level.

### 3.4 A case study: RiboRL provides biological insights for translation elongation

Given the clear association between the RiboRL prediction and the underlying biological rules, we further assessed the ability of our framework to provide biological insights in a specific scenario. In particular, we attempted to analyze the signals provided by RiboRL in transmembrane (TM) protein biogenesis. To facilitate the experiment, we downloaded 839 yeast transmembrane proteins from the UniProt database [45] and extracted their last transmembrane domains that are flanked by at least 10 amino acids at both ends. Then we aligned all the transmembrane sequences with respect to the start positions of their transmembrane domains, and calculated, for each position, the average predicted ribosome density as well as the frequency of being selected as a rationale by the RiboRL model (Figure 4). Intriguingly, we observed a clear difference between the transmembrane domains (with a length of around 20 amino acids) and the flanking water soluble regions. Consistent with previous reports [46, 47], we first observed a significant decrease in ribosome density within the transmembrane domain, presumably due to the increase in translation speed in the structured protein regions (Figure 4). In addition, we also noticed that the codons in transmembrane regions are less likely to be selected as rationales by the PolicyNet compared to the water soluble regions (*P <* 10^*−*300^ by two-sided Mann-Whitney U test), indicating a relatively less predictive power of these features in predicting local ribosome density (Figure 4). As the transmembrane domains are typically composed of continuous hydrophobic amino acids, this result suggested that the flanking hydrophilic amino regions potentially make more contributions to the regulation of translation elongation dynamics near the transmembrane domains.

**Figure 4:**
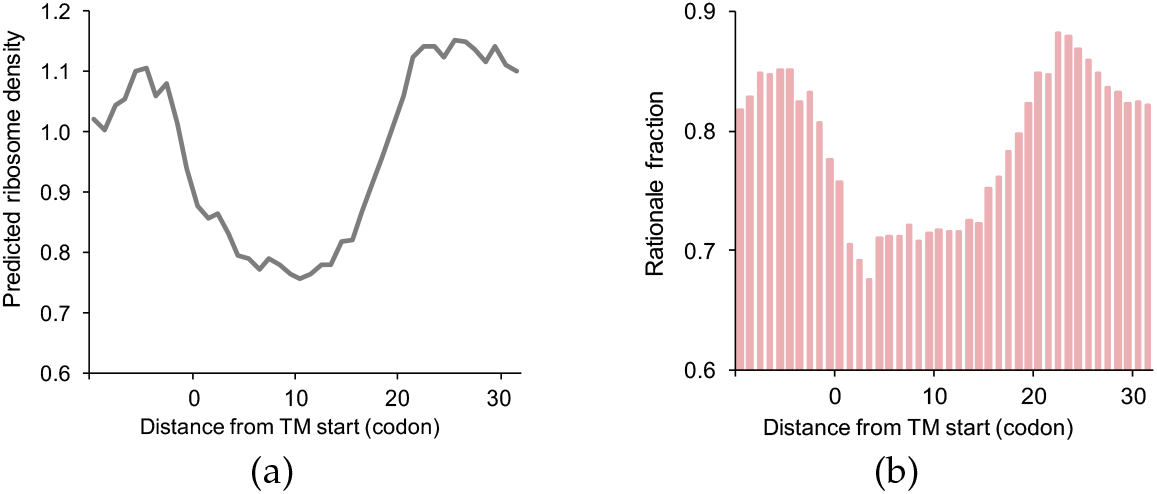
A case study of applying RiboRL to generate biological insights for translation elongation dynamics in transmembrane (TM) protein biogenesis. (a) The average predicted ribosome density and (b) the fractions of being chosen as rationales at different positions with respect to the start positions of the transmembrane domains whose lengths are about 20 amino acids.

## 4 Discussion and conclusions

Understanding the sequence features that contribute to translation elongation dynamics has long been regarded as an important biological problem. On the other hand, the recent development of deep learning methods have already demonstrated their promising application in deciphering complex biological systems [48–53]. In this work, empowered by the deep reinforcement learning strategy, we propose a context-dependent feature selection approach to obtain the most relevant sequence features within the input sequence of each sample, thus providing a more accurate and robust prediction for ribosome density on the mRNA transcripts.

Admittedly, the studies of technical bias and shortcomings [43, 54] of ribosome profiling have also attracted wide interest in the computational biology community [55–58]. Without any doubt, understanding these problems would also be beneficial for deciphering the biological determinants of translation elongation dynamics. Nevertheless, the recent experimental efforts have demonstrated that the computational models based on the current conventional ribosome profiling pipelines can provide sufficient guidance for the design of translation output [19].

The current study serves as the *first* attempt to apply reinforcement learning to study the translation elongation process. In the future, we will extend the reward function in our framework to further explore the possibility of generating the rationales with specific properties, e.g., positively or negatively contributing rationales to provide more detailed insights into the translation elongation process. In addition, we expect our framework would be useful in the identification of key sequence elements for protein design [59] and synthetic biology [60] in the future.

## Acknowledgments

This work was supported in part by the National Natural Science Foundation of China (61472205, 81630103, 61472205, 81630103 and 61836004) and China’s Youth 1000-Talent Program, and Beijing Brain Science Special Project (No.Z181100001518006).

## Footnotes

1 In reinforcement learning, the reward of an action is defined by the summation of the final reward for the action sequence and the corresponding intermediate reward for each step. Since RiboRL does not define intermediate reward, the returned reward of an action is equal to the final reward.

## References

[1] N. Ingolia, “Ribosome footprint profiling of translation throughout the genome,” Cell, vol. 165, no. 1, pp. 22–33, 2016.

[2] T. E. Dever, J. D. Dinman, and R. Green, “Translation elongation and recoding in eukaryotes,” Cold Spring Harbor perspectives in biology, p. a032649, 2018.

[3] C. Kimchi-Sarfaty, J. M. Oh, I.-W. Kim, Z. E. Sauna, A. M. Calcagno, S. V. Ambudkar, and M. M. Gottesman, “A“ silent” polymorphism in the MDR1 gene changes substrate specificity,” Science, vol. 315, no. 5811, pp. 525–528, 2007.

[4] R. Ishimura, G. Nagy, I. Dotu, H. Zhou, X.-L. Yang, P. Schimmel, S. Senju, Y. Nishimura, J. H. Chuang, and S. L. Ackerman, “Ribosome stalling induced by mutation of a CNS-specific tRNA causes neurodegeneration,” Science, vol. 345, no. 6195, pp. 455–459, 2014.

[5] G. A. Brar, “Beyond the triplet code: context cues transform translation,” Cell, vol. 167, no. 7, pp. 1681–1692, 2016.

[6] T. Tuller, Y. Y. Waldman, M. Kupiec, and E. Ruppin, “Translation efficiency is determined by both codon bias and folding energy,” Proceedings of the National Academy of Sciences, vol. 107, no. 8, pp. 3645–3650, 2010.

[7] M. Stadler and A. Fire, “Wobble base-pairing slows in vivo translation elongation in metazoans,” RNA, vol. 17, no. 12, pp. 2063–2073, 2011.

[8] W. Qian, J.-R. Yang, N. M. Pearson, C. Maclean, and J. Zhang, “Balanced codon usage optimizes eukaryotic translational efficiency,” PLoS genetics, vol. 8, no. 3, p. e1002603, 2012.

[9] C. A. Charneski and L. D. Hurst, “Positively charged residues are the major determinants of ribosomal velocity,” PLoS Biol, vol. 11, pp. 1–20, Mar. 2013.

[10] A. Dana and T. Tuller, “The effect of tRNA levels on decoding times of mRNA codons,” Nucleic Acids Research, 2014.

[11] C. Pop, S. Rouskin, N. T. Ingolia, L. Han, E. M. Phizicky, J. S. Weissman, and D. Koller, “Causal signals between codon bias, mRNA structure, and the efficiency of translation and elongation,” Molecular Systems Biology, vol. 10, no. 12, 2014.

[12] D. E. Weinberg, P. Shah, S. W. Eichhorn, J. A. Hussmann, J. B. Plotkin, and D. P. Bartel, “Improved ribosome-footprint and mRNA measurements provide insights into dynamics and regulation of yeast translation,” Cell Reports, vol. 14, no. 7, pp. 1787–1799, 2016.

[13] S. Zhang, H. Hu, J. Zhou, X. He, T. Jiang, and J. Zeng, “Analysis of ribosome stalling and translation elongation dynamics by deep learning,” Cell systems, vol. 5, no. 3, pp. 212–220, 2017.

[14] K. D. Duc and Y. S. Song, “The impact of ribosomal interference, codon usage, and exit tunnel interactions on translation elongation rate variation,” PLoS genetics, vol. 14, no. 1, p. e1007166, 2018.

[15] N. T. Ingolia, S. Ghaemmaghami, J. R. S. Newman, and J. S. Weissman, “Genome-wide analysis *in vivo* of translation with nucleotide resolution using ribosome profiling,” Science, vol. 324, no. 5924, pp. 218–223, 2009.

[16] N. T. Ingolia, J. A. Hussmann, and J. S. Weissman, “Ribosome profiling: Global views of translation,” Cold Spring Harbor perspectives in biology, p. a032698, 2018.

[17] T.-Y. Liu and Y. S. Song, “Prediction of ribosome footprint profile shapes from transcript sequences,” Bioinformatics, vol. 32, no. 12, pp. i183–i191, 2016.

[18] P. B. F. O’Connor, D. E. Andreev, and P. V. Baranov, “Comparative survey of the relative impact of mRNA features on local ribosome profiling read density,” Nature Communications, vol. 7, pp. 12915–,Oct. 2016.

[19] R. J. Tunney, N. J. McGlincy, M. E. Graham, N. Naddaf, L. Pachter, and L. Lareau, “Accurate design of translational output by a neural network model of ribosome distribution,” Nature Structure & Molecular Biology, 2018.

[20] R. Tibshirani, “Regression shrinkage and selection via the lasso,” Journal of the Royal Statistical Society. Series B (Methodological), pp. 267–288, 1996.

[21] B. Langmead and S. L. Salzberg, “Fast gapped-read alignment with bowtie 2,” Nature methods, vol. 9, no. 4, p. 357, 2012.

[22] M. Pertea, D. Kim, G. M. Pertea, J. T. Leek, and S. L. Salzberg, “Transcript-level expression analysis of RNA-seq experiments with hisat, stringtie and ballgown,” Nature protocols, vol. 11, no. 9, pp. 1650–1667, 2016.

[23] D. Kim, B. Langmead, and S. L. Salzberg, “Hisat: a fast spliced aligner with low memory requirements,” Nature methods, vol. 12, no. 4, p. 357, 2015.

[24] T. Lei, R. Barzilay, and T. Jaakkola, “Rationalizing neural predictions,” in EMNLP, pp. 107–117, 2016.

[25] D. Bahdanau, K. Cho, and Y. Bengio, “Neural machine translation by jointly learning to align and translate,” arXiv preprint arXiv:1409.0473, 2014.

[26] A. Graves, A. rahman Mohamed, and G. E. Hinton, “Speech recognition with deep recurrent neural networks,” In Acoustics, speech and signal processing (icassp), 2013 IEEE international conference on. IEEE, pp. 6645–6649, 2013.

[27] A. G. Barto, R. S. Sutton, and C. W. Anderson, “Neuronlike adaptive elements that can solve difficult learning control problems,” IEEE transactions on systems, man, and cybernetics, no. 5, pp. 834–846, 1983.

[28] K. Cho, B. van Merrienboer, C. Gulcehre, F. Bougares, H. Schwenk, and Y. Bengio, “Learning phrase representations using RNN encoder-decoder for statistical machine translation,” EMNLP, p. 1724–1734, 2014.

[29] R. S. Sutton, A. G. Barto, et al., Reinforcement learning: An introduction. MIT press, 1998.

[30] V. Mnih, K. Kavukcuoglu, D. Silver, A. A. Rusu, J. Veness, M. G. Bellemare, A. Graves, M. Riedmiller, A. K. Fidjeland, G. Ostrovski, et al., “Human-level control through deep reinforcement learning,” Nature, vol. 518, no. 7540, p. 529, 2015.

[31] A. Conneau, D. Kiela, H. Schwenk, L. Barrault, and A. Bordes, “Supervised learning of universal sentence representations from natural language inference data,” in Proceedings of the 2017 Conference on Empirical Methods in Natural Language Processing, pp. 670–680, 2017.

[32] L. Mou, Z. Lu, H. Li, and Z. Jin, “Coupling distributed and symbolic execution for natural language queries,” in ICML, pp. 2518–2526, 2017.

[33] R. J. Williams, “Simple statistical gradient-following algorithms for connectionist reinforcement learning,” Machine Learning, vol. 8, pp. 229–256, 1992.

[34] X. Liu, L. Mou, H. Cui, Z. Lu, and S. Song, “Jumper: Learning when to make classification decisions in reading,” in IJCAI, 2018.

[35] Q. L. J. P. Yang Liu, Prajit Ramachandran, “Stein variational policy gradient,” in UAI, 2017.

[36] Y. Kim, “Convolutional neural networks for sentence classification,” in EMNLP, pp. 1746–1751, 2014.

[37] M. Sundararajan, A. Taly, and Q. Yan, “Axiomatic attribution for deep networks,” in ICML, 2017.

[38] A. Shrikumar, P. Greenside, and A. Kundaje, “Learning important features through propagating activation differences,” in ICML, 2017.

[39] R. Singh, J. Lanchantin, A. Sekhon, and Y. Qi, “Attend and predict: Understanding gene regulation by selective attention on chromatin,” in Advances in Neural Information Processing Systems, pp. 6788–6798, 2017.

[40] H. Hu, A. Xiao, S. Zhang, Y. Li, X. Shi, T. Jiang, L. Zhang, L. Zhang, and J. Zeng, “Deephint: Understanding HIV-1 integration via deep learning with attention,” Bioinformatics, 2018.

[41] Y. Luo, J. Ma, Y. Liu, Q. Ye, T. Ideker, and J. Peng, “Deciphering signaling specificity with interpretable deep neural networks,” in Research in Computational Molecular Biology: 22nd Annual Conference, RECOMB 2018, Paris, France, April 21-24, 2018, Proceedings, Springer International Publishing, 2018.

[42] S. J. Sheather and M. C. Jones, “A reliable data-based bandwidth selection method for kernel density estimation,” Journal of the Royal Statistical Society. Series B (Methodological), vol. 53, no. 3, pp. 683–690, 1991.

[43] C. G. Artieri and H. B. Fraser, “Accounting for biases in riboprofiling data indicates a major role for proline in stalling translation,” Genome Research, 2014.

[44] P. M. Sharp and W.-H. Li, “The codon adaptation index – a measure of directional synonymous codon usage bias, and its potential applications,” Nucleic Acids Research, vol. 15, no. 3, pp. 1281–1295, 1987.

[45] U. Consortium et al., “Uniprot: the universal protein knowledgebase,” Nucleic Acids Research, vol. 46, no. 5, p. 2699, 2018.

[46] R. Saunders and C. M. Deane, “Synonymous codon usage influences the local protein structure observed,” Nucleic Acids Research, vol. 38, no. 19, pp. 6719–6728, 2010.

[47] S. Pechmann and J. Frydman, “Evolutionary conservation of codon optimality reveals hidden signatures of cotranslational folding,” Nat Struct Mol Biol, vol. 20, pp. 237–243, Feb. 2013.

[48] B. Alipanahi, A. Delong, M. T. Weirauch, and B. J. Frey, “Predicting the sequence specificities of DNA-and RNA-binding proteins by deep learning,” Nat Biotech, vol. 33, pp. 831–838, Aug. 2015.

[49] J. Zhou and O. G. Troyanskaya, “Predicting effects of noncoding variants with deep learning-based sequence model,” Nat Meth, vol. 12, pp. 931–934, Oct. 2015.

[50] S. Zhang, H. Hu, T. Jiang, L. Zhang, and J. Zeng, “TITER: predicting translation initiation sites by deep learning,” Bioinformatics, vol. 33, no. 14, pp. i234–i242, 2017.

[51] N. H. Tran, X. Zhang, L. Xin, B. Shan, and M. Li, “De novo peptide sequencing by deep learning,” Proceedings of the National Academy of Sciences, vol. 114, no. 31, pp. 8247–8252, 2017.

[52] R. Zhang, Y. Wang, Y. Yang, Y. Zhang, and J. Ma, “Predicting ctcf-mediated chromatin loops using ctcf-mp,” Bioinformatics, vol. 34, no. 13, pp. i133–i141, 2018.

[53] Y. Y. Lu, J. Lv, Y. Fan, and W. S. Noble, “DeepPINK: reproducible feature selection in deep neural networks,” in NIPS, 2018.

[54] J. A. Hussmann, S. Patchett, A. Johnson, S. Sawyer, and W. H. Press, “Understanding biases in ribosome profiling experiments reveals signatures of translation dynamics in yeast,” PLoS Genet, vol. 11, pp. 1–25, Dec. 2015.

[55] H. Wang, J. McManus, and C. Kingsford, “Accurate recovery of ribosome positions reveals slow translation of wobble-pairing codons in yeast,” in Research in Computational Molecular Biology: 20th Annual Conference, RECOMB 2016, Santa Monica, CA, USA, April 17-21, 2016, Proceedings (M. Singh, ed.), pp. 37–52, Cham: Springer International Publishing, 2016.

[56] H. Fang, Y.-F. Huang, A. Radhakrishnan, A. Siepel, G. J. Lyon, and M. C. Schatz, “Scikit-ribo enables accurate estimation and robust modeling of translation dynamics at codon resolution,” Cell systems, vol. 6, no. 2, pp. 180–191, 2018.

[57] D. Zhao, W. Baez, K. Fredrick, and R. Bundschuh, “Riboprop: A probabilistic ribosome positioning algorithm for ribosome profiling,” Bioinformatics, 2018.

[58] F. Lauria, T. Tebaldi, P. Bernabò, E. J. Groen, T. H. Gillingwater, and G. Viero, “ribowaltz: Optimization of ribosome p-site positioning in ribosome profiling data,” PLoS Computational Biology, vol. 14, no. 8, p. e1006169, 2018.

[59] E. Natan, T. Endoh, L. Haim-Vilmovsky, T. Flock, G. Chalancon, J. T. Hopper, B. Kintses, P. Horvath, L. Daruka, G. Fekete, et al., “Cotranslational protein assembly imposes evolutionary constraints on homomeric proteins,” Nature structural & molecular biology, vol. 25, no. 3, p. 279, 2018.

[60] P. Ciryam, R. I. Morimoto, M. Vendruscolo, C. M. Dobson, and E. P. O’Brien, “In vivo translation rates can substantially delay the cotranslational folding of the escherichia coli cytosolic proteome,” Proceedings of the National Academy of Sciences, vol. 110, no. 2, pp. E132–E140, 2013.

